# An electromyography-based multi-muscle fatigue model to investigate operational task performance

**DOI:** 10.1101/2023.11.15.567264

**Authors:** Leonardo H. Wei, Suman K. Chowdhury

**Affiliations:** Industrial Manufacturing and Systems Engineering, Texas Tech University, Lubbock, United States

**Keywords:** multiple muscle fatigue model, static and dynamic exertions, electromyography, shoulder joint fatigue

## Abstract

We developed a multi-muscle fatigue model (MMFM) by incorporating electromyography (EMG)-based amplitude and frequency parameters, the fast-to-slow twitch muscle fiber ratio, a time multiplier to linearize the cumulative effect of time, and a muscle multiplier to standardize the combined effect of the number of muscles being considered. We validated the model by investigating fatigue development patterns of ten male subjects performing one sustained-till-exhaustion static and two repetitive dynamic tasks (low and high task difficulty levels) using 0.91 Kg and 2.72 Kg dumbbells. The results indicated that the MMFM was sensitive to fatigue-related neuromuscular changes and predicted shoulder joint fatigue accurately.

## Introduction

Muscle fatigue has been defined as the reduction of force-generating capacity of the muscular system, usually seen as a failure to maintain or to develop expected muscle force or power (Vøllestad 1997). It is a complex physiological state of a muscle, primarily developed during prolonged voluntary muscle contractions through two mechanisms: 1) the accumulation of metabolites (e.g., lactic acid) within muscle fibers and 2) a reduction in the brain motor drives (Enoka and Duchateau 2008). Many previous studies have observed a substantial impact of muscle fatigue on physical performance (Lyons et al. 2006; Dupuis et al. 2022; Tornero-Aguilera et al. 2022), cognitive performance (Martin et al. 2020), and cognitive alertness (Martin et al. 2020) of human subjects in various operational environments. In addition, several studies have used muscle fatigue as the biomarker to identify the risk of musculoskeletal (MSK) injuries in occupational settings (Anton et al. 2001; Dubowsky et al. 2008; Nimbarte et al. 2013; Luger et al. 2016; Rashedi and Nussbaum 2016), sports (Mueller-Wohlfahrt et al. 2013; Goes et al. 2020) and daily activities (Shan et al. 2013; Legan and Zupan 2022). As MSK injuries are prevalent across occupational settings (40% of all MSK work-related injuries (BLS 2020), sports (76% of all injuries among 567 athletes (Goes et al. 2020)), and daily activities (39.6% out of 535 students (Legan and Zupan 2022)), an accurate assessment of muscle fatigue is essential to determine appropriate risk mitigation strategies in order to reduce the risk of MSK injury across all operational environments.

In previous studies, researchers have mainly used the following five assessment methods to quantify muscle fatigue: 1) changes in the maximum voluntary contraction (MVC) (Vøllestad 1997), 2) changes in the endurance time (Liu B et al. 2018), 3) changes in metabolite concentration (Jebelli et al. 2020), 4) near-infrared spectroscopy (Scano et al. 2020), and 5) electromyography (Cifrek et al. 2009) and 6) visual analog scale (Borg 1982). Among these methods, surface electromyography (SEMG) is a preferred assessment method due to its high precision, non-invasiveness, and unobtrusiveness (Merletti et al. 2001; Bandpei et al. 2014). Several different SEMG signal processing techniques—such as amplitude and frequency parameters—were used to understand the onset of muscle fatigue state in various operational activities (Luttmann et al. 2000; Jebelli and Lee 2019; Liu S-H et al. 2019). Among them, the most commonly used techniques to understand muscle fatigue include: 1) an increase in the time domain parameters i.e., integrated EMG (IEMG), root mean square (RMS) and mean absolute value (MAV), and normalized mutual information (NMI) (Kawczyński et al. 2015) and 2) a decrease in the spectral domain parameters i.e., median frequency (MDF) and the mean frequency (MPF) (Mathur et al. 2005; Bosch T et al. 2007; Calder et al. 2008). This is because the onset of muscle fatigue causes a shift in the recruitment of fast (high-twitch), high-fatiguing fibers to a large amount of slow (low-twitch), low-fatiguing fibers in order to compensate for the external force requirement. As a result, the EMG signal pattern shows an increase in amplitude and a decrease in frequency power with the manifestation of muscle fatigue. However, there are putative evidence for these aforementioned EMG-based fatigue assessment methods (Farina and Enoka 2011). For example, some isometric studies have found an expected trend of increase in amplitude and decrease in frequency with the development of fatigue (Bosch T et al. 2007; Nussbaum 2008; Tucker et al. 2009). In contrast, some isometric (Clancy et al. 2008; □ström et al. 2009) and dynamic studies found these spectral- and amplitude-based fatigue measures to be insensitive to muscle fatigue (Sood et al. 2007; Bosch Tim et al. 2011; Bosch Tim et al. 2012). These putative evidences on the efficacy of EMG-based measures was mainly due to the fact that the fatigue development and progression are affected by the task intensity and duration and the muscles being involved (Basmaijan and De Luca 1985).

In an attempt to correctly assess muscle fatigue, several previous studies developed empirical and theoretical models for a better understanding of muscle fatigue. The empirical muscle fatigue models were validated by either SEMG and/or force data. For example, Ma et al. (2009) developed a dynamic fatigue model to predict muscle fatigue considering external load and personal contributions (maximum voluntary contraction). Their formulation had two first-order differential equations – one for the fatigue development process and the other one for the recovery process. The authors validated their model using SEMG and force data and proclaimed that the model requires further experimental validation for more dynamic work scenarios. On the contrary, the theoretical muscle fatigue models were based on the mathematical representation of the physiological muscle process during prolonged muscle contraction and were mainly presumed by existing evidence. For instance, a dynamic muscle model to describe activation, fatigue, and recovery levels of muscle fibers was proposed by Liu JZ et al. (2002). The model described the muscle force induced over a given activation-fatigue-recovery time frame, relating the input of the brain, assumed constant, to the force. Similarly, Frey-Law et al. (2012) proposed a biophysical model based on the active, fatigued, and resting muscle states to describe the optimal parameters of fatigue and recovery. This model was later modified due to the over-prediction of fatigue during complex conditions by Looft et al. (2018), who included a rest recovery parameter to represent muscle recovery better. Potvin and Fuglevand (2017) also proposed a fatigue model based on the fatigue of motor units. They simulated motor unit firing rates and isometric forces of a 120-motor unit muscle. However, their model is limited to isometric muscle contraction scenarios. In addition, it does not consider the recovery stage from fatigue; this is important because the degree of muscle fatigue depends not only on prior muscle activity but also on the muscle resting period. Despite the existence of aforementioned fatigue models to assess individual muscles’ physiological state, a multi-muscle fatigue assessment model is essential to evaluate operational activities because the performance of a task requires the coordinated activations of multiple muscles and joint(s). Thus, the fatigue assessment of only one or two muscles may not represent a precise fatigue estimation of multiple muscle groups involved during a task performance. Moreover, fatigue development and progression of a muscle depends not only on its’ own fatigue state and brain motor drive but also on the synergistic activation and fatigue level of other muscles involved in performing the same task (Kouzaki and Shinohara 2006; Szucs et al. 2009).

To our knowledge, only one previous study proposed a multi-muscle fatigue index using EMG-based amplitude and frequency parameters to evaluate shoulder muscle fatigue (McDonald et al. 2019). They considered the summation of all fatigued muscles’ scaled amplitude and frequency values. The scaling multipliers were calculated based on the maximal correlation between their formulation and perceived fatigue level. Moreover, the number of muscles contributing to the total summation of their fatigue score was multiplied as a linearization term to consider the effects of muscle state in the overall fatigue. However, each muscle’s anatomical characteristics were not considered in the formulation. As the EMG amplitude and spectrum depend on muscle anatomical characterizations (e.g., cross-sectional area, muscle fiber composition) (Farina and Holobar 2016), a multi-muscle fatigue model without the consideration of muscle anatomical characterizations may lead to erroneous muscle fatigue assessment. For example, muscles with a higher percentage of slow twitch fibers have higher fatigue resistance capacity compared to muscles with a higher percentage of fast twitch fibers (type II) (Scott et al. 2001). Consequently, a multi-muscle fatigue model based on the muscle fiber characteristics is expected to facilitate more accurate measurement of the physiological strain of a joint.

Therefore, this study aimed to develop a multi-muscle fatigue model based on the SEMG data and individual muscles’ anatomical characterization. Our developed multi-muscle fatigue index was expected to facilitate both fatigue level and fatigue type of individual muscle groups for any given task. To validate the multi-muscle fatigue model, sub-maximal static and repetitive dynamic exertions of the shoulder joint were considered under different muscle loading conditions. We hypothesized that physically demanding tasks would lead to greater muscle fatigue than physically less-demanding tasks. Thus, a multi-muscle fatigue model is expected to facilitate a precise estimation of joint fatigue and reduce the risks of the incidences of musculoskeletal disorders (MSDs).

## Materials and Methods

### Multi-Muscle Fatigue Model Development

The mathematical formulation of our proposed multi-muscle fatigue model (MMFM) considered the instantaneous contributions of all activated and fatigued muscles for a given task, the type of fatigue (localized peripheral muscle fatigue versus brain or cognitive central fatigue), anatomical characteristics (muscle fiber compositions), and scaling multipliers to linearize the formulation. The EMG-based joint analysis of spectral and amplitude (JASA) technique was the basis for identifying the type of fatigue and the number of fatigued muscles (i.e., muscle state) (Luttmann et al. 1996). Several previous studies implemented the JASA technique to explore the insights of muscle state for various operational activities (Luttmann et al. 2000; Lin et al. 2004; Dos Santos et al. 2017; Ding et al. 2020). Briefly, the JASA method, as shown in Figure1, describes four possible muscle state scenarios based on the shifts in EMG amplitude and frequency values: 1) the increase in both EMG amplitude (*A*) and EMG frequency (*f*) values indicate an increase in muscle force (1^st^ quadrant), 2) the decrease in *f* but an increase in *A* refers to the state of muscle fatigue (2^nd^ quadrant), 3) the decrease of *f* and *A* indicates the brain central fatigue, i.e., decrease in muscle force due to the decline in brain effort (3^rd^ quadrant), and 4) the increase *f* of but an increase a decrease in *A* refers to the state of muscle recovery (4^th^ quadrant), i.e., the recovery of brain and muscle from the fatigue state. We used this EMG-based JASA method to identify the muscle states, the types of fatigue (localized peripheral muscle fatigue and brain/cognitive central fatigue) a muscle undergoes, and the total number of fatigued muscles.

**Figure 1:**
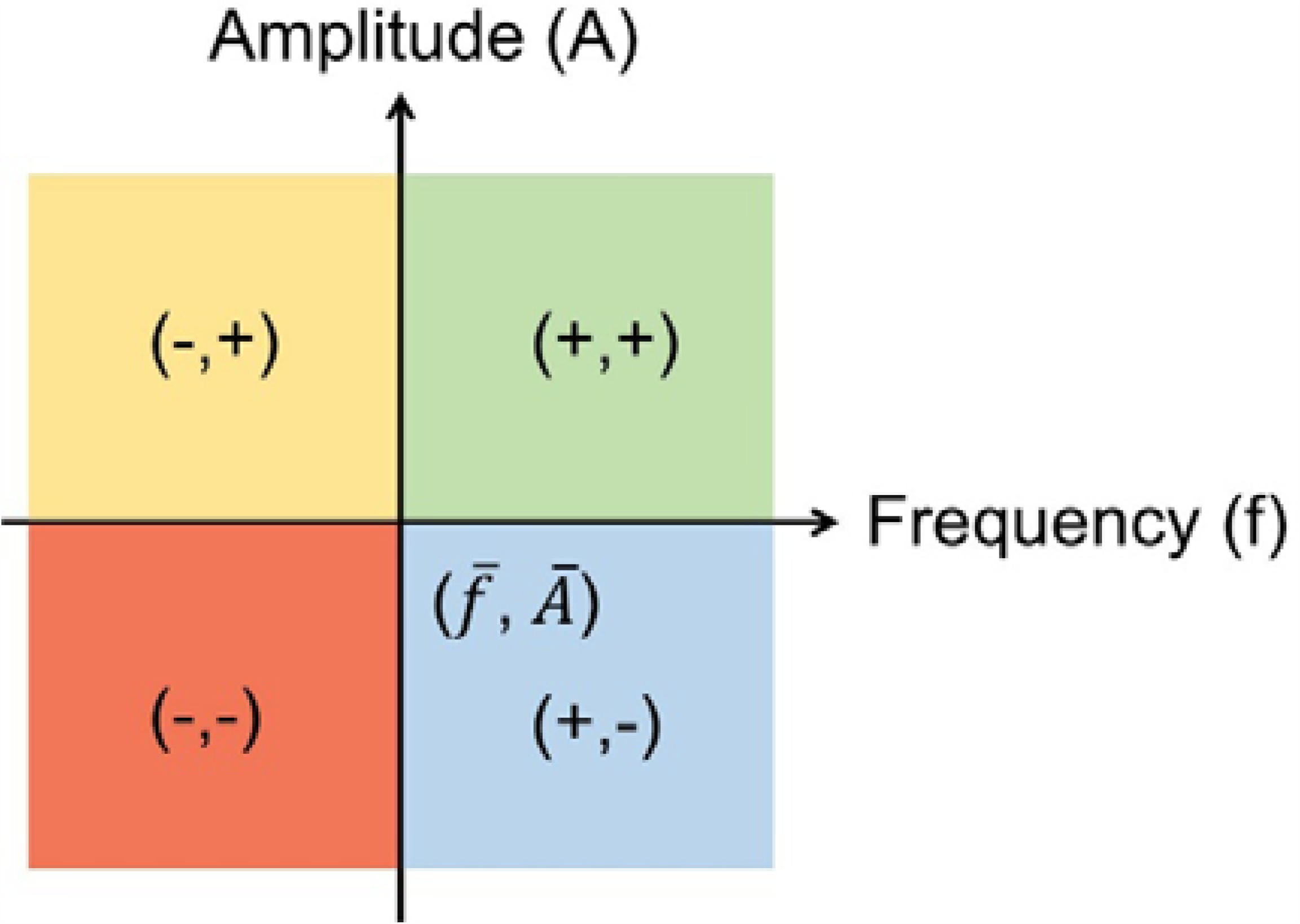
A schematic presentation of the Joint Analysis of Spectral and Amplitude (JASA) to understand electromyography (EMG)-based muscle states. The symbols, 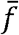 and *Ā* refer to average EMG amplitude and median frequency of baseline non-fatigued EMG signals, respectively. Both plus (+) and minus (-) signs respectively indicate an increase and a decrease in instantaneous EMG amplitude and/or frequency values.

Furthermore, muscle co-activation, task duration, the amount of fatigue at any given time (time dependency), and the proportion of fiber types affect the progression of muscle fatigue. For example, the higher a task is sustained (task duration), the more metabolite substrates accumulate in muscle fibers, which, in turn, influence the capability of myosin and actin filaments to form cross-bridges (Fitts 2008; Debold 2012). If the task is continued without a rest period, the fatiguing fibers cannot get enough time to recover. That means muscle fatigue can be considered cumulative at any given time step (time dependency). In addition, the proportion of fast to slow twitch fibers affects the firing rate (frequency and amplitude) of each muscle (Fitts 2006; Bogdanis 2012). Moreover, depending on how muscles fatigue over the course of task performance, the central nervous system changes its’ its muscle co-activation strategy (both patterns and combination), i.e., it can target different muscles to co-activate in order to optimize energy expenditure and/or even to improve task accuracy (Dul et al. 1984; Missenard et al. 2008).

Therefore, we used the following criteria to develop our proposed MMFM for any given fatiguing static and/or dynamic tasks: (1) muscles with a higher proportion of type II fibers fatigue faster, (2) an increase in the SEMG amplitude and a decrease in the SEMG frequency is the biomarker of localized peripheral muscle fatigue, (3) a decrease in both SEMG amplitude and frequency is the biomarker of cognitive central fatigue, i.e., a decrease in brain effort to generate the required force, (4) tasks that engage a higher number of muscles to have the ability to sustain longer, i.e., such task have the flexibility to engage alternate muscles if a group of muscles is fatigued, and (5) muscles that fatigued faster contribute to a faster rate of physiological strain on the joint. By combining these five criteria, we formulated the following mathematical model of MMFM:

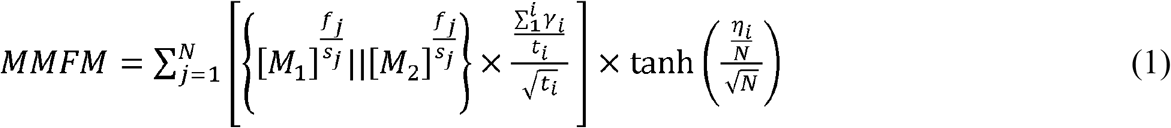

where, *i* is an index for time step, *j* is an index for the number of muscles, *N* is the total number of muscles for a given task, *η* _*i*_ is the number of fatigued muscles at any given time, *t*_*i*_ is the total number of time steps (windows) at *i*^*th*^ time step, *γ* _*i*_ is a counter for the number of time steps (windows) a muscle experiences both localized peripheral and cognitive central fatigue by *i*^*th*^ time step, 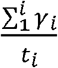 is a time domain multiplier to account for the number of times a muscle experience fatigue by time step *i*,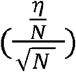 is a muscle multiplier,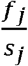 is the ratio between fast (*f*_*j*_) and slow twitch (*S*_*j*_) fibers of a muscle (*j*), *A*_*i*_ and *F*_*i*_ are the amplitude and frequency at i^th^ time step, and 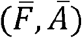 represents baseline frequency and amplitude, which are the average of the first three-time steps. *M* 1and *M* 2 respectively represent localized peripheral muscle (*j*) fatigue and cognitive central fatigue at time step (*i*) :

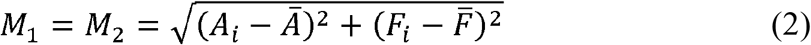

We used two normalized reference values: 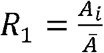 and 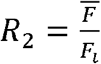 and three following decision criteria to determine if a muscle undergoes localized peripheral muscle fatigue or cognitive/brain central fatigue, or no fatigue for any given time step, *i*.

- Criterion 1: *R*_1_ ≥1and *R*_2_ ≥1, then *M* 1 is considered.
- Criterion 2: *R*_1_ ≤1and *R*_2_ ≥1, then *M* 2 is considered.
- Criterion 3: *R*_2_ <1, then *M* 1= *M* 2=0.

The first term of the MMFM formulation, 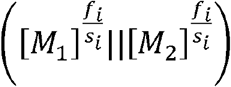, indicates the type of fatigue a muscle undergoes. The mathematical operator, “OR” symbol (||), was used in the first term to reflect that either cognitive central or physical fatigue can occur at any given time step. The fast-to-slow twitch fiber ratio 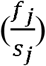 was used as a power exponent in the first term to emphasize the manifestation of fatigue in muscles having a greater amount of fast-twitch fibers than slow-twitch fibers, as they are more vulnerable to fatigue-related muscle injuries. The second term 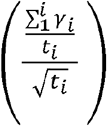, a time-domain multiplier, was used to linearize the non-linearity of the first term, which is due mainly to the cumulative effect of fatigue over the course of time. The second term also standardizes the cumulative fatigue assessment across tasks regardless of their durations. Lastly, we combined the products of first and second terms of individual muscles across all time steps in order to estimate the overall joint fatigue (i.e., the fatiguing performance of all muscles) for a given task. Thus, the last term 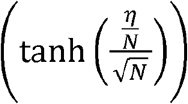, a muscle multiplier was a hyperbolic function and its’ its values range between 0 and 1. This hyperbolic function was used to further normalize the summation of the product of the first and second terms regardless of the number of muscles being considered in those two terms.

### Model Validation

#### Experiment Design Participants

To test the appropriateness of the MMFM, we recruited ten healthy male participants (age = 28.50 ± 3.55 years; weight = 74.9 ± 7.76 Kg; height = 175.9 ± 3.78 cm) to perform fatiguing exertions under both static and dynamic conditions. All participants were required to be free from any type of musculoskeletal, degenerative, or neurological disorders and have no history of neck, back, and shoulder injury or notable pain. Participants who met the inclusion criteria were asked to read and sign a consent form approved by the local Institutional Review Board (IRB # 1505685703) before participating in the experimental tasks.

#### Experimental procedure

The static exertion involved holding a weight with the right arm at shoulder flexion of 90° till exhaustion with no elbow flexion and shoulder abduction. The dynamic exertion involved repetitive exertions from elbow height to mid-upper arm height (the mid-point between the elbow and shoulder heights). The starting and ending point of the exertions were 30% and 100% of thumb-tip reach at a mid-sagittal plane, respectively. Both static and dynamic exertions were performed using two different dumbbells of 0.91 and 2.72 Kg in a custom-made workstation, as shown in Figure 2. There was a total of six experimental conditions, in which each condition was repeated three times. All tasks were randomly selected for each participant, and their muscle activity was recorded using EMG sensors placed in eight different shoulder muscles: medial deltoid, anterior deltoid, posterior deltoid, supraspinatus, infraspinatus, teres major, biceps, and triceps.

**Figure 2:**
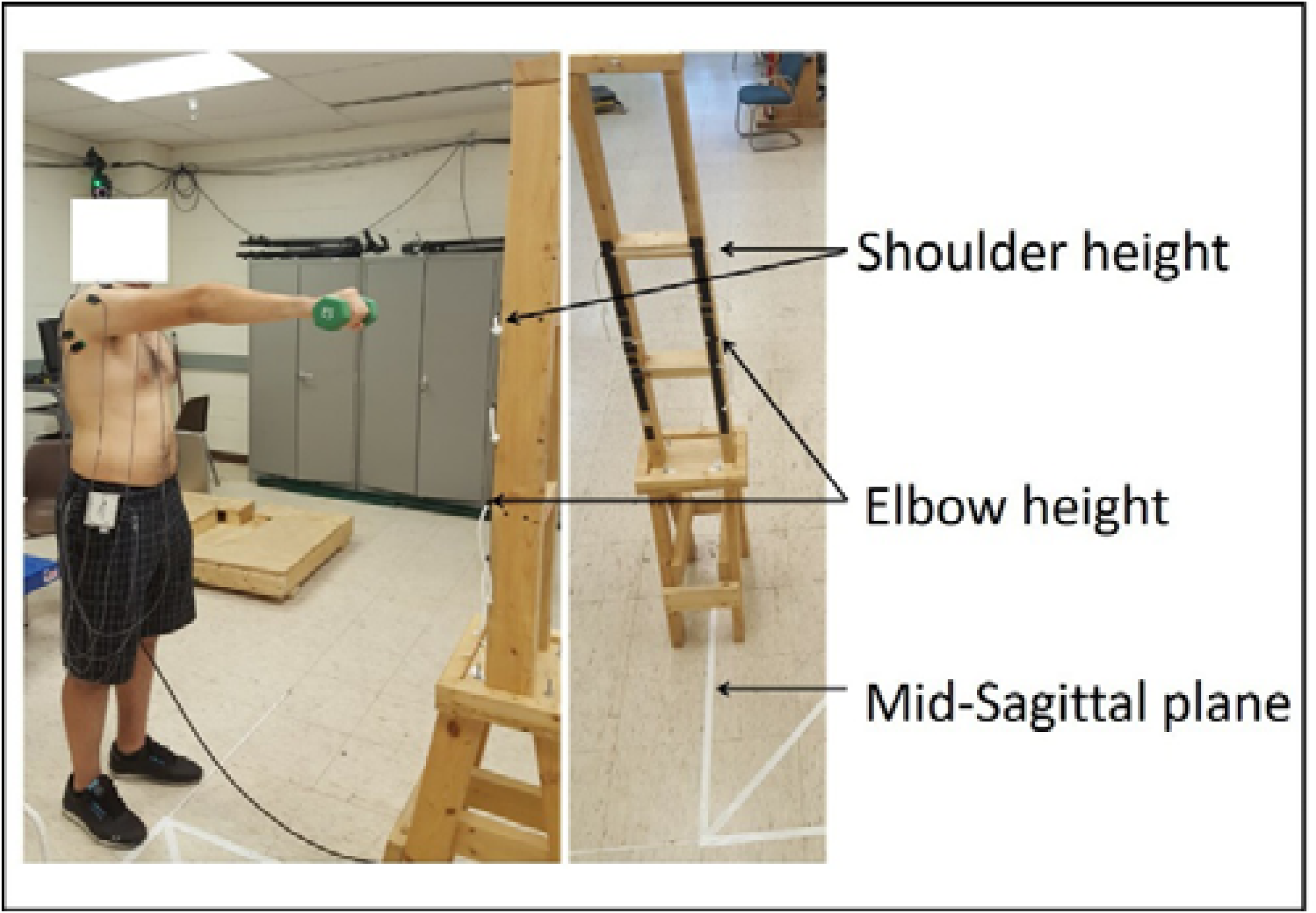
(a) Surface electromyography (EMG) apparatus (right) and Biodynamic apparatus (left) used in this study to respectively measure individual muscles’ activity and maximum voluntary contraction.

#### Data Acquisitions

A Bagnoli-16 desktop SEMG system (Delsys Inc., Boston, USA) was used to collect the muscle activation data from the shoulder muscles (Figure 3). The EMG sensors used for the data acquisition are parallel bar single differential surface electrodes with an inter-electrode distance of 10 mm, common mode rejection ratio of 92 dB, input impedance greater than 1 MΩ, and a sampling rate of 2000 Hz. The maximum voluntary contractions (MVC) of each muscle were performed at an isokinetic dynamometry (HUMAC NORM, Computer Sports Medicine (CSMi), Stoughton, MA) (Figure 3).

**Figure 3:**
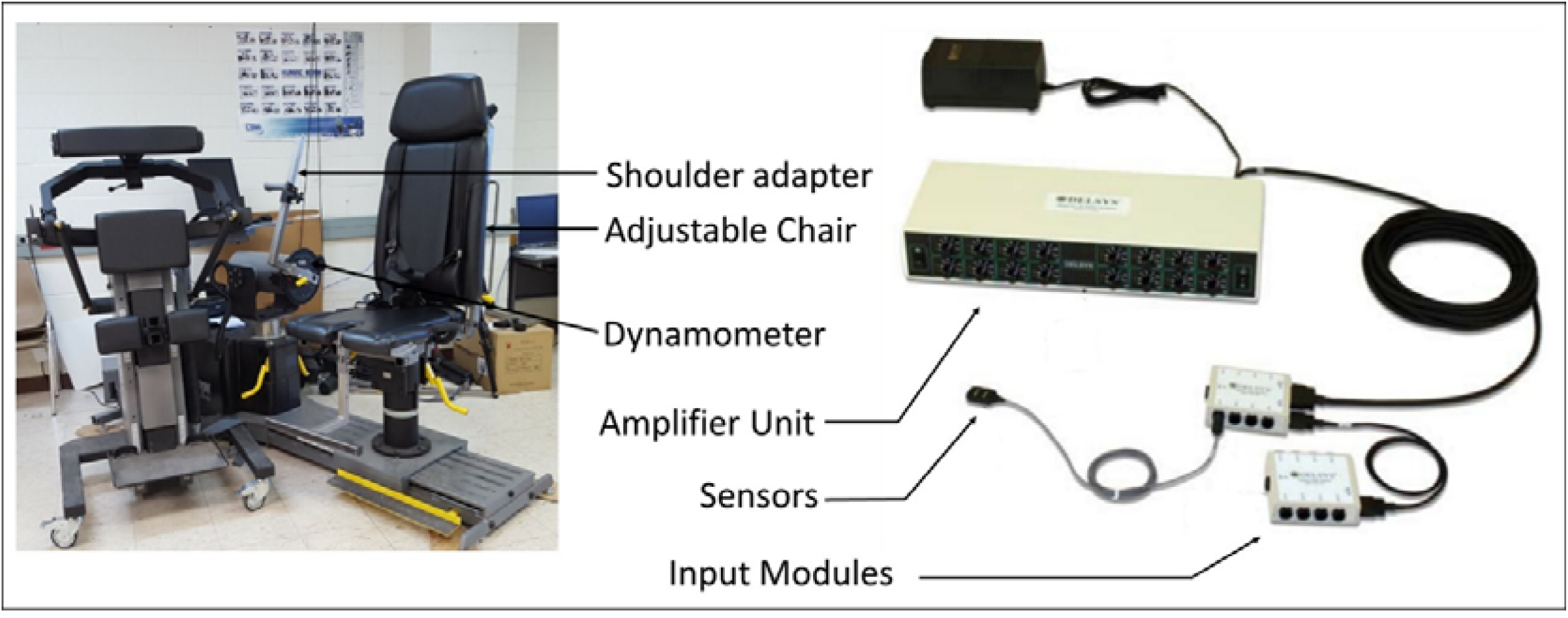
A custom-designed experimental setup to simulate the experimental tasks.

Prior to the experimental tasks, participants were asked to perform two consecutive MVCs for each muscle, with a rest period of at least two minutes between them to reduce the effects of fatigue. After the MVC data collection, participants were permitted to elapse a rest period of ten minutes prior to beginning the static and dynamic tasks. During dynamic exertions, participants were allowed to choose the pace of the repetitive exertions. However, they were asked to maintain the same pace throughout the experiment. The pace was controlled using a metronome. A rest period of 3 minutes was provided between the tasks. SEMG data were recorded continuously during the tasks. At the beginning and at the end of each task, the participants were asked to rate their self-perceived shoulder discomfort on a 0 – 10 Borg CR-10 scale (Borg 1982).

#### Data Analysis

The filtered signals were full-wave rectified and smoothed using an eight-order Butterworth low-pass filter. The SEMG signals of each muscle were normalized with respect to the peak of their corresponding MVC signal to minimize between-subject or between-muscle errors. The IEMG was calculated for the muscle contraction duration (*τ*_*i*_) and the mean absolute value for each muscle was calculated using Equation 3:

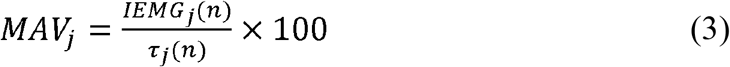

where *MAV*_*j*_, *τ*_*i*_, and *IEMG*_*i*_ are respectively mean absolute value of muscle *j* at *n*^*th*^ exertion, contraction duration for the *j*^*th*^ muscle at *n*^*th*^ nexertion, and the integrated EMG for *j*^*th*^ muscle at *n*^*th*^ exertion.

During repetitive dynamic exertions, a period of muscle contraction is characteristically followed by a period of muscle relaxation when the muscle is returned to its original resting position. The total duration of each muscle contraction was about 4 s throughout any given task. Therefore, a 4-s long muscle contraction period (*τ*_*i*_) was chosen for all trials and all subjects (Figure 4). In order to calculate the MMF value of a repetitive exertion, we slid a 500 ms (with overlapping of 250 ms window) moving window throughout the 4-s contraction frame and calculated MF and MAV for each moving window (Figure 4), which were subsequently used as inputs to the JASA decision criteria (Figure 1). M1 and M2 fatigue components were estimated if any instantaneous MAV and MF of a 500-ms window exceeded baseline non-fatigued amplitude (*Ā*) and frequency 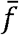 values. These baseline values for dynamic and static exertion signals were calculated by averaging MAV and MF values of the first three repetitions of dynamic exertion signals and first three window sizes of static exertion signals, respectively.

**Figure 4:**
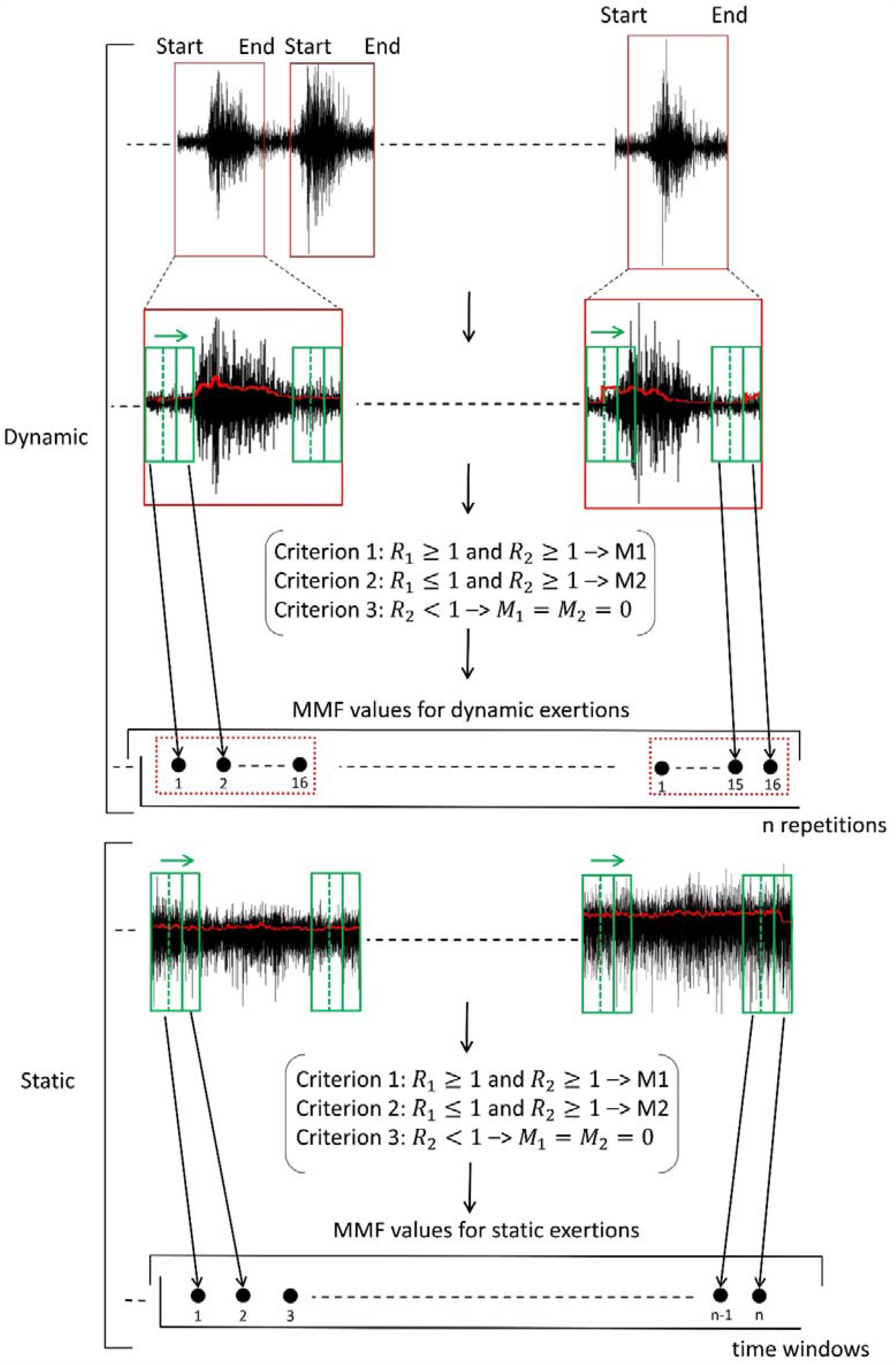
A schematic presentation of the electromyography (EMG) data processing workflow for static and dynamic exertion signals. Red boxes in dynamic signals represent the contraction frame of individual repetitive exertions. A moving window of 500 ms with a 250 ms overlapping window (both are green color-coded) was slid in each dynamic contraction frame, resulting in 16 different moving windows (shown as solid red dots). Similar moving windows were also used for static exertion signals. R1 and R2 are the decision criteria to determine the physical (M1) and cognitive/brain central (M2) fatigue components of individual muscles for the MMF estimation.

Then M1 and M2 component of each muscle was powered to their fast-to-slow twitch fiber ratio (Table 1) in order to incorporate the time-variant muscle fiber dynamics as per the muscle size principle (Fling et al. 2009). Then, the MMF value of repetitive exertion was estimated by averaging the MMF values of all 16 windows of a contraction frame. Similarly, SEMG signals of static trials were rectified and smoothed, and the MMF values were estimated for each 500 ms (with overlapping of 250 ms window) moving window of the complete trial (Figure 4). In addition, MMF values were calculated for all three repetitions of six experimental task conditions. Moreover, individual subjects’ perceived effort ratings (Borg scale) were averaged for each experimental condition in order to correlate and validate their corresponding objective fatigue (MMFM) values.

**Table 1.**
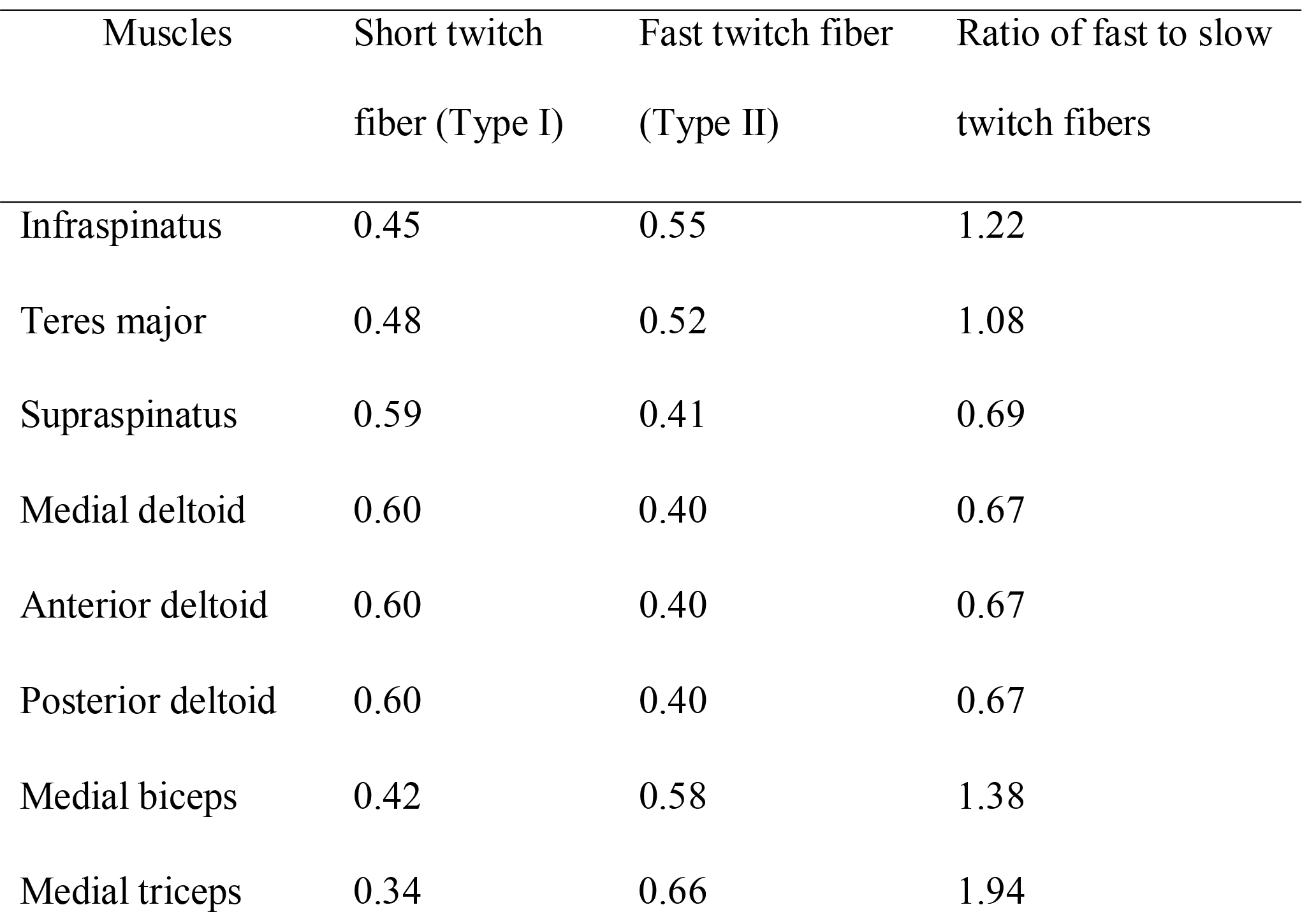
Fiber type compositio ns in the shoulder muscles (Karlsson 1992).

#### Statistical Analysis

The descriptive statistics (mean and standard error) of MMF values for all repetitive exertions of an experimental condition were calculated by averaging them across all repetitions and all subjects. In order to validate our MMFM, we investigated linear trends (slopes) of MMF values over time for all six experimental conditions. In addition, we investigated physical (M1), cognitive central (M2), or no-fatigue muscle states for the effects of task difficulty level (low vs. high) on dynamic exertion tasks and load conditions (0.91 Kg vs. 2.72 Kg) on both static and dynamic exertion tasks by using one-way analysis of variance test with 95% confidence level. The dependent variable was percent of fatigue contribution 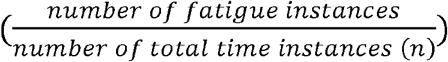 and the independent variable was fatigue type (M1, M2, and no-fatigue conditions). A post hoc Tukey’s statistical analysis was also employed if the levels of each experimental condition had significantly contributed to M1, M2, and no-fatigue muscle states. Furthermore, we investigated M1, M2, and no-fatigue states of individual muscles and their contributions to the total MMF values. The MMFM was also validated by investigating linear trend (slope) and Pearson correlation coefficient between subjective (Borg’s perceived rating) and objective (MMF values) fatigue values for each experimental condition.

## Results

### MMF Trends

The MMF values of 2.72-Kg and 2-Kg static exertions were notably higher than those of 2.72-Kg and 0.91-Kg dynamic exertions across all time steps (Figure 5). The 2.72-Kg static exertion displayed the steepest slope trend (m = 0.91) and the highest peak MMF value of 19.3 among all trials (task difficulty level). Both peak and slope values of 2.72-Kg static exertion were about 30.6% and 7.1% greater than those (m = 0.85; peak = 14.78) of the 0.91-Kg static exertion, respectively. Both peak and slope values of 0.91-Kg and 2.72-Kg dynamic exertions showed smaller peak and slope values than their static counterparts. For instance, the peak and slope values of 2.72-Kg-high dynamic exertions (m = 0.43; peak = 7.16) were respectively 62.9% and 52.7% lesser than those of 2.72-Kg static exertions (m = 0.91; peak = 19.30), whereas, the peak and slope values of 0.91-Kg-high dynamic exertions (m = 0.13; peak = 3.75) were respectively 74.6% and 84.7% lesser compared to those of 0.91-Kg static exertions. Likewise, 0.91-Kg-low (m = 0.05; peak = 3.11) and 2.72-Kg-low (m = 0.08; peak = 3.83) dynamic exertions displayed 94.1% and 78.9% lower peak values and 91.2% and 80.2% lower slope values compared to those of 0.91-Kg and 2.72-Kg static exertions, respectively. The peak values across all task difficulty levels occurred at the last timestep (100% of total time) (Figure 5).

**Figure 5:**
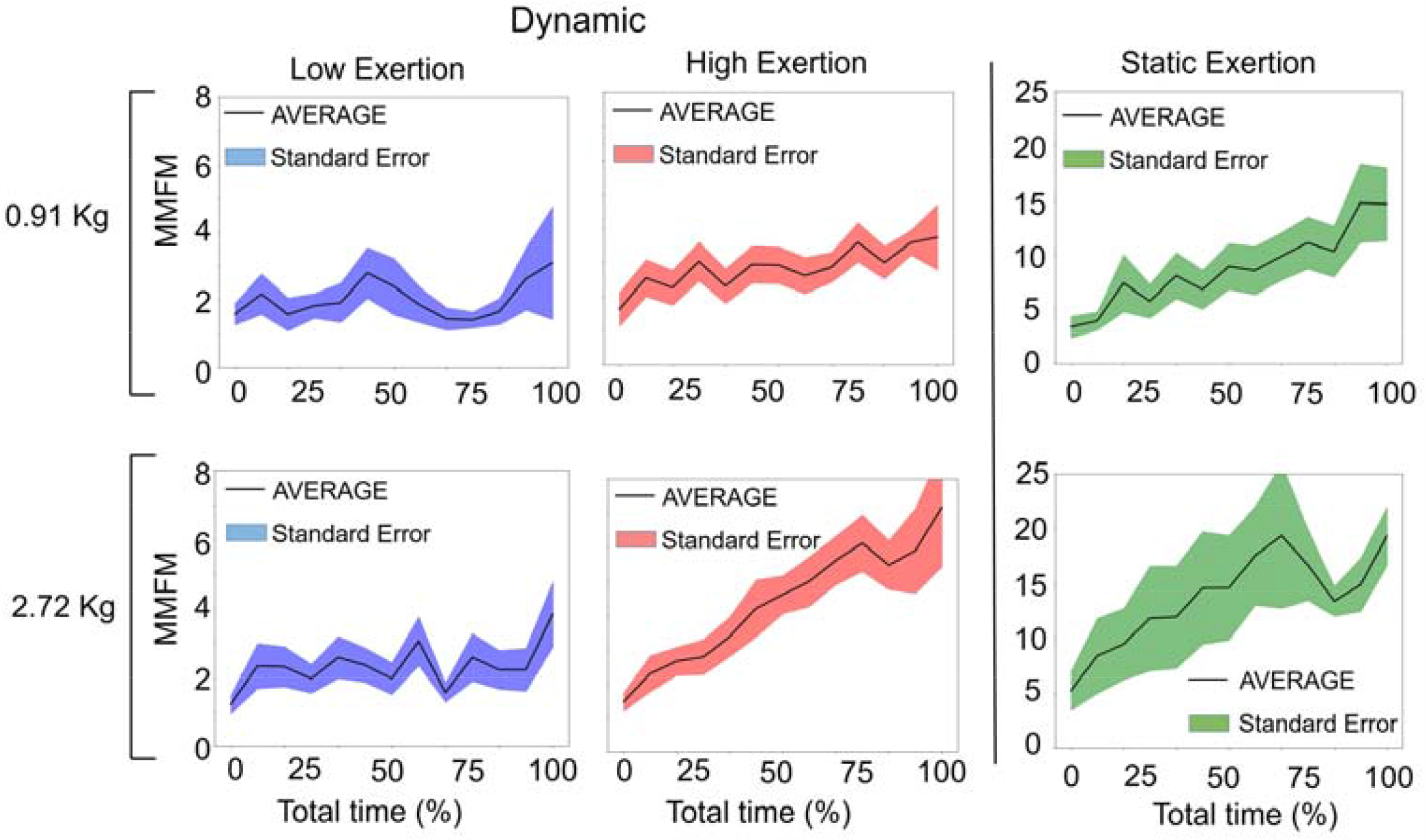
The MMFM trends for each of the six experimental conditions: two different load conditions (0.91 and 2.72 Kg) and three task difficulty levels (low and high dynamic and static exertions). Solid lines represent overall fatigue trends, whereas shaded regions indicate corresponding standard errors for each of the experimental conditions.

The muscle state data retrieved from the MMFM showed that showed that the quantity of physical fatigue (M1) was significantly (p-value <0.01) higher than the central fatigue (M2) across all task difficulty levels for most task duration shoulder muscles significantly (p-value <0.01) experienced physical fatigue (M1) than brain fatigue (M2) across all task difficulty levels for most of the task duration (62.8% of total), except for the 0.91-Kg-low dynamic exertion, where M1 was insignificantly (p-value>0.24) greater than M2 (Figure 6). The 2.72-Kg static exertion displayed the greatest proportion of M1 (∼ 65% of 1024) and M2 (∼ 25% of 1024) states and the least proportion of no-fatigue (∼10%) state, followed by 2.72-Kg-high dynamic, 0.91-Kg-static, 2.72-Kg-low dynamic, 0.91-Kg-high dynamic, and 0.91-Kg-low dynamic exertion tasks. In contrast, low dynamic exertion tasks exhibited no-fatigue muscle state for significantly (0.91 Kg: p-value<0.0001, 2.72 Kg: p-value<0.0001) a higher proportion of task duration (46% of total) compared to M1 and M2 states. Likewise, we compared the individual muscle state data (as shown in Figure 7) in order to understand their contributions to total joint fatigue (MMF values). We observed that triceps, middle, anterior, and posterior deltoid muscle groups, rotator cuff muscle groups (supraspinatus, teres major, and infraspinatus muscles) experienced a higher proportion of M1 state for most of the task duration compared to M2 and no-fatigue states for both static exertion tasks (Figure 7). They contributed a higher percentage of M1 state, followed by M2 state, to MMF values (MMFM data), particularly at 75 – 100% of the task duration. Interestingly, the anterior deltoid and teres major muscles showed a greater proportion of M2 fatigue state than M1 and no-fatigue state for 0.91-Kg static exertion tasks. During low dynamic exertion tasks (for both 0.91 Kg and 2.72 Kg load conditions), anterior deltoid, infraspinatus, and triceps muscles primarily contributed to M1 and M2 fatigue states, specifically, at 50 – 100% of the task duration (Figure 7), whereas, the rest of the muscles showed a predominant amount of no-fatigue state throughout entire task duration (Figure 7).

**Figure 6:**
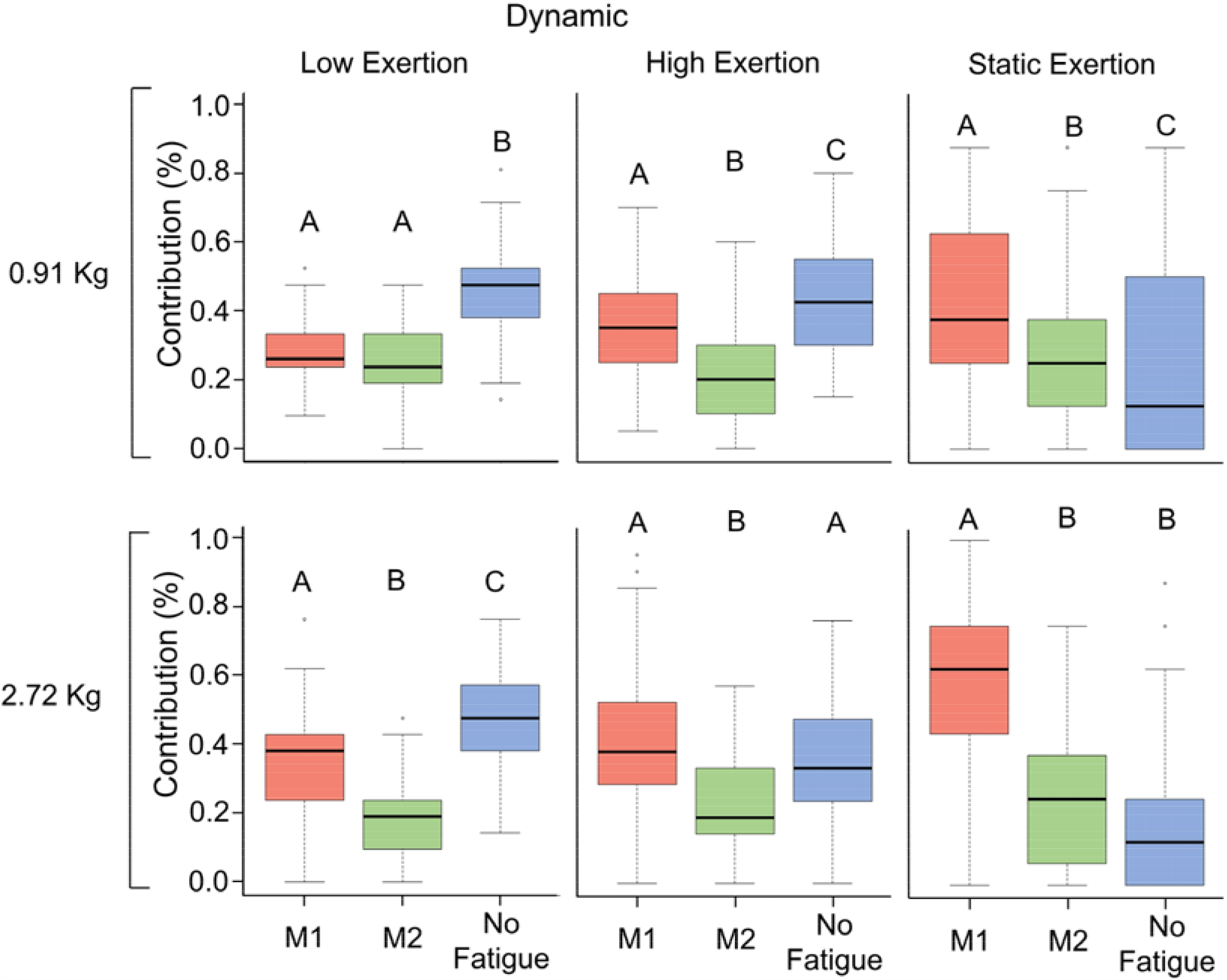
Task assessment based on the proportion of physical fatigue (M1), brain/cognitive central fatigue (M2), and no-fatigue states for all muscles (joint) retrieved from the multi-muscle fatigue model data. Symbols A, B, and C represent post-hoc differences between the levels of each experimental condition. A different letter between the levels indicates a statistically significant difference.

**Figure 7:**
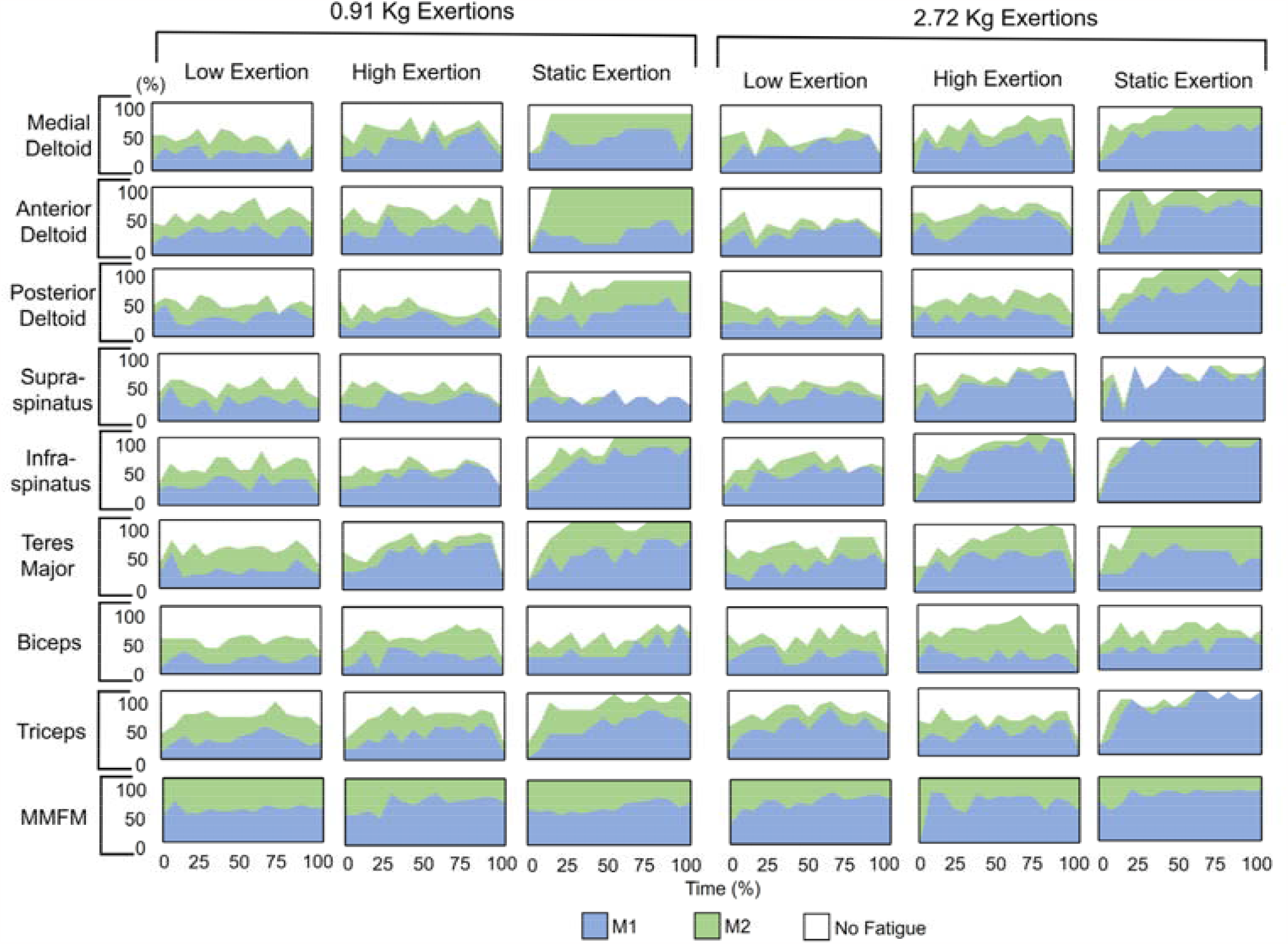
Proportions of physical fatigue (M1), brain/cognitive central fatigue (M2), and no-fatigue states for the individual muscles and their contributions to the overall MMF values (last row) for each load condition and task difficulty levels over time. Blue, green, and white-coded regions represent M1, M2, and no-fatigue states, respectively.

### Subjective fatigue and MMF values

The MMF values for each task difficulty level were compared with their respective Borg’s perceived exertion rating data (subjective fatigue), and their relationship for individual load conditions and position (low vs. high exertion) are respectively presented in Figures 8 and 9. Figure 8 displayed a greater decreasing trend (m= -0.29) and a higher increasing trend (m = 0.28) for 0.91-Kg and 2.72-Kg dynamic tasks, respectively, indicating that the majority of the participants perceived no-to-low amount of effort (0 – 4; with a mean and standard deviation of 1.75 ± 1.08) for dynamic tasks using 0.91-Kg load condition, whereas they perceived a greater amount of medium-to-high effort (4 – 8; with a mean and standard deviation of 4.33 ± 2.23) for dynamic tasks using 2.72 Kg load condition. Similarly, Figure 9 exhibited a slight downward (negative) slope (m = -0.02) for low dynamic exertion tasks, whereas a greater amount of upward (positive) trend (m = 0.14) was observed for high dynamic exertion tasks, indicating that subjects perceived a low sense of effort (1 – 4; with a mean and standard deviation of 2.3 ± 1.60) for low dynamic exertions and a medium-to-high sense of effort (3 – 8; with a mean and standard deviation of 3.78 ± 2.40) for high dynamic exertions.

**Figure 8:**
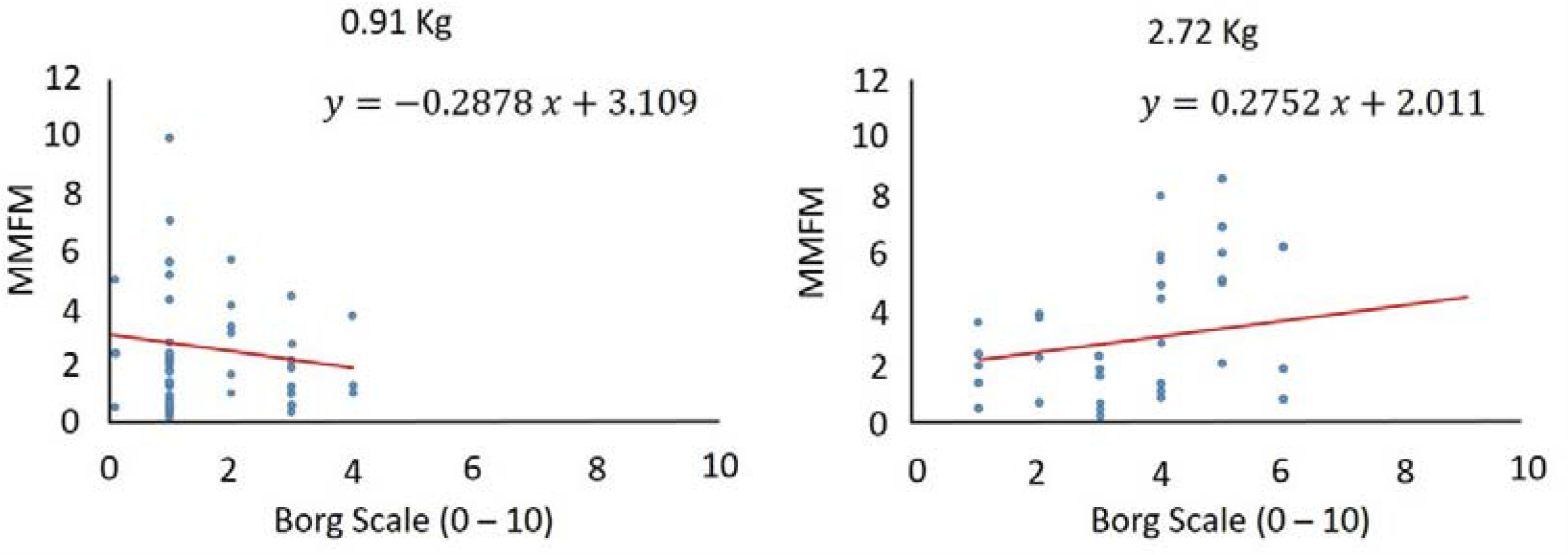
Comparison between Borg’s perceived rating (X-axis) and MMF value (Y-axis) for 0.91 kg and 2.72 kg dynamic tasks (averaged across all task difficulty levels).

**Figure 9:**
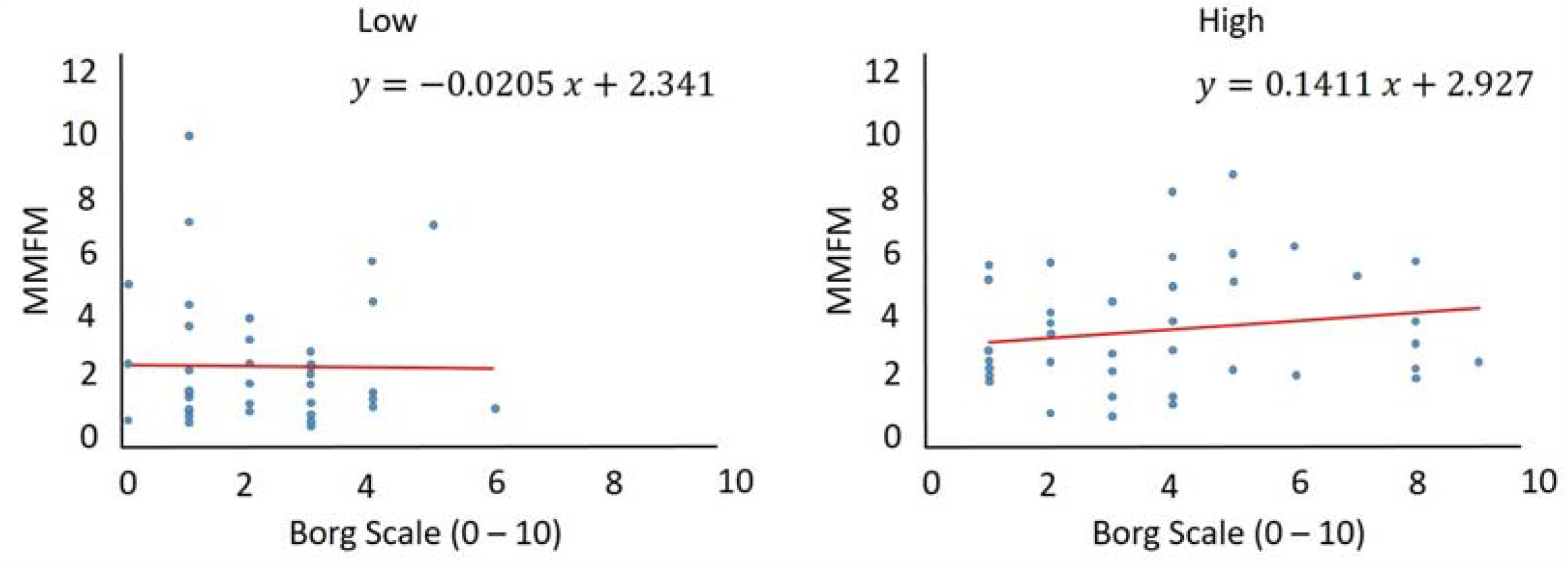
Comparison between Borg’s perceived rating (X-axis) and MMF values (Y-axis) for high and low dynamic exertion tasks (averaged across all load conditions).

In addition, the perceived exertion data for 0.91 Kg and 2.72 Kg static tasks were found to be in the range of an effort level between 4 – 7 (with a mean and standard deviation of 6.1 ± 1.2) and 7 – 10 (with a mean and standard deviation of 8.6 ± 2.3), respectively.

## Discussion

In this study, we developed a mathematical formulation of MMFM and validated the formulation via a laboratory-based human subject experiment, in which participants performed fatiguing static and dynamic shoulder exertions using two load conditions – 0.91 and 2.72 Kg. Using the SEMG data collected from eight shoulder muscles, MMF values of the shoulder joint were calculated and validated against Borg’s perceived exertion data (subjective fatigue) for each experimental condition. Results exhibited higher objective (MMF values) and subjective (perceived exertion scores) fatigue values with the increase in the load and task difficulty levels. Moreover, static exertion tasks resulted in higher fatigue (both objective and subjective) than dynamic exertion tasks.

Our MMFM displayed considerably higher joint fatigue (MMF values) for static exertion tasks compared to dynamic exertion tasks for both load conditions, suggesting that static tasks are comparatively more intense and demanding. This can be explained by the steady-state muscle contraction (with no rest periods) characteristics, which makes blood vessels of contracted muscles remain partially restricted and inhibits the oxygen supply and the removal of lactic acid (Enoka 1995; Chen and Lee 1998; Korshøj et al. 2016; Luger et al. 2016). In contrast, during dynamic exertions, muscles undergo relaxation periods between contractions and intermittently let blood vessels carry oxygen and remove lactic acid from the muscles. Consequently, muscles showed significantly a greater amount of physical (muscle) fatigue (M1) than cognitive central fatigue (M2) and no-fatigue states during static exertion tasks than dynamic exertion tasks.

The unique characteristics of our multi-muscle fatigue (i.e., joint fatigue) formulation were that it considered both brain/cognitive central fatigue (M2) and localized peripheral muscle fatigue (M1) of individual muscles of a joint. Such EMG-based fatigue formulation could open the door to understanding time- and muscle-specific onsets of muscle and brain central fatigue experienced by the individual muscles during operational task performance. Our results showed that physical fatigue (M1) was found to be higher with the increase in task difficulty level (from 0.91 Kg to 2.72 Kg load and low to high dynamic task conditions), indicating that a higher amount of muscle fibers were engaged to compensate the increased metabolic energy requirement due to the increased force (load effect) and moment-arm (task difficulty effect) demands on the shoulder joint. On the contrary, the increase in cognitive central fatigue (M2) proportion was primarily found in the third and fourth quarters of the task duration, suggesting the decrease in brain/cognitive central effort due to the cumulative manifestation of localized peripheral muscle fatigue in the 2^nd^ quarter. The static exertion tasks showed this time-and task-dependent dynamics of M1 and M2 fatigue manifestations more prominently. These findings based on our MMFM formulation assured the force-fatigability relationship, in which a greater weight/force exertion triggers faster and higher fatigue levels (Lind and Petrofsky 1979; Bellemare and Grassino 1982; Bigland□Ritchie and Woods 1984; Clark and Carter 1985).

The subjective fatigue data also showed that the perceived effort level increased with the increase in load and task difficulty levels. The perceived effort data for tasks using 0.91 Kg load condition and low exertion level displayed a nominal relationship (no-to-low amount of effort) with their corresponding objective fatigue data (MMF values). In contrast, the perceived effort data for tasks using the 2.72 Kg load condition and high exertion level exhibited an increasing trend and relationship (medium-to-high amount of effort) with their corresponding objective fatigue data (MMF values). Our finding agreed with the findings of some previous studies, in which they evaluated the effect of exercise-induced task difficulty on their mental/cognitive loading and found that with the increase in exercise difficulty, participants experienced a higher level of cognitive demand (Cian et al. 2001). In summary, Borg’s perceived rating data validated the capability of MMFM in characterizing and predicting muscle and brain central fatigue levels precisely for various operational tasks.

Previous EMG-based fatigue models primarily used amplitude and/or frequency parameters to identify fatigue (Mathur et al. 2005; Bosch T et al. 2007; Calder et al. 2008), with no consideration of the dynamics of brain effort (Ma et al. 2009) and the time-dependent manifestation of muscle fatigue (McDonald et al. 2019), though central brain (or central) fatigue has been found to affect the voluntary drive of muscles (Contessa et al. 2016). Furthermore, the longer a muscle contracts, the more fatigue accumulates in muscles (Hogan et al. 1998; Enoka and Duchateau 2008). On the other hand, our MMFM formulation accounted for both time and brain effects on individual muscles. Nevertheless, our MMF formulation employed time and muscle multipliers to linearize the cumulative summation of physical and cognitive central fatigue components over time and the number of muscles being considered for a given task. The muscle multiplier standardized the fatigue contributions regardless of the number of muscles that experience fatigue at any given time. Similarly, the time multiplier standardized the cumulative summation of prior fatigue states of individual muscles at each time period. In addition, the MMFM formulation accounted fast-to-slow twitch fiber ratio to prioritize muscles that are fast-fatiguing and more vulnerable to injury.

Though the MMFM was observed to be sensitive to fatigue-related neuromuscular changes, there are a few study limitations that need to be acknowledged. First, the MMFM was validated for a limited number of static and dynamic exertion tasks, which were performed in standard neutral standing posture with a fixed speed and duration. The tasks in occupational settings are not always performed under such conditions. Secondly, the participant pool was limited to university-aged students with little to no manual materials handling experience. Experienced workers may exhibit different material handling and muscle recruitment strategies. Third, only male participants were recruited. However, the manifestation of fatigue in female participants may follow a different pattern. Fourth, our MMFM formulation did not explicitly consider muscle co-contraction dynamics and muscle physiological cross-sectional area, which could have provided a slightly different fatigue assessment, particularly during dynamic exertion tasks. Given that our goal was to formulate a multi-muscle fatigue model based on EMG signal patterns and muscle anatomical characterization and validate the model for various operational tasks, the experimental tasks and participant pool included in this study may be adequate. Nonetheless, future studies should assess the model with a wide range of participant pools and various operational tasks. In conclusion, the present study established a fatigue model that can be useful to assess and monitor both physical (muscle) and cognitive central (brain) fatigue levels of multiple muscles simultaneously as well as their joined combined fatigue level for various operational tasks., and thus help Thus, the model is believed to help practitioners to determine mitigation strategies to reduce the risk of MSDs in the operational environment.

## Acknowledgments

The authors acknowledge Anindita Das for her contribution in data processing.

